## Supplementary File for "Social-affective features drive human representations of observed actions"

**a.** Breakdown of scene setting and number of agents across the two final stimulus sets.

| | Scene setting | | | Number of agents | | | |
|---|---|---|---|---|---|---|---|
| | Indoors | Outdoors | Unclear | 0 | 1 | 2 | 3+ |
| Video set 1 | 89 | 60 | 3 | 8 | 52 | 50 | 42 |
| Video set 2 | 39 | 26 | 0 | 0 | 21 | 29 | 15 |

**b.** Features quantified in both stimulus sets and used to generate feature RDMs in the representational similarity analysis. *This feature was generated as a binary RDM clustering videos by category.

| Feature name | Feature type | Computation method | Numeric representation (per video) |
|---|---|---|---|
| Pixel value | Visual | Automatic extraction | Real-valued vector (480,000D) |
| Hue | Visual | Automatic extraction | Real-valued vector (480,000D) |
| Saturation | Visual | Automatic extraction | Real-valued vector (480,000D) |
| Watermark | Visual | Experimenter labels | Value (0-1) |
| Gist | Visual | Automatic extraction | Real-valued vector (512D) |
| Environment | Visual | Experimenter labels | Value (0-1) |
| Optic flow | Visual | Automatic extraction | Real-valued vector (480,000D) |
| AlexNet Conv1 | Visual | Automatic extraction | Real-valued vector (69,984D) |
| AlexNet FC8 | Visual | Automatic extraction | Real-valued vector (1000D) |
| Action category | Action | Experimenter labels | Binary RDM* |
| Effectors | Action | Experimenter labels | Binary vector (5D) |
| Transitivity | Action | Behavioral ratings | Value (1-5) |
| Activity | Action | Behavioral ratings | Value (1-5) |
| Valence | Social-affective | Behavioral ratings | Value (1-5) |
| Arousal | Social-affective | Behavioral ratings | Value (1-5) |
| Sociality | Social-affective | Behavioral ratings | Value (1-5) |
| Number of agents | Social-affective | Experimenter labels | Value (0-3) |
